# Catalytic promiscuity potentiated the divergence of new cytochrome P450 enzyme functions in cyanogenic defense metabolism

**DOI:** 10.1101/398503

**Authors:** Brenden Barco, Lara Zipperer, Nicole K. Clay

**Author notes:** Correspondence should be addressed to B. Barco.

## Abstract

Cytochrome P450 monooxygenases (P450s) constitute the largest metabolic enzyme family in plants, responsible for synthesizing hundreds of thousands of specialized metabolites with essential roles in chemical defenses against herbivores and pathogens (Banks et al., 2011; Nelson and Werck-Reichhart, 2011; Wurtzel and Kutchan, 2016). Substrate promiscuity has been documented to play a central role in the evolution of plant specialized metabolic enzymes (Weng et al., 2012; Leong and Last, 2017), however most plant P450s are highly substrate-specific (Verpoorte, 2013). Here, we show the rapid inversion of primary and weak secondary (promiscuous) catalytic activities between two distinct yet evolutionarily linked multifunctional P450s, CYP71A12 and CYP71A13, based on intramolecular epistasis of two amino acid residues under positive selection in CYP71A12. Furthermore, we uncover previously undocumented catalytic activity during the inversion as well as naturally occurring amino acid substitution patterns that could have been present in evolutionary intermediates between the two enzymes. Comparative expression profiling and homology modeling reveal that natural selection acted on the promoter of CYP71A13 and the substrate-recognition elements of CYP71A12 to improve the efficiencies of their promiscuous reactions. The rise in catalytic promiscuity potentiated the divergence of new P450 enzyme functions in cyanogenic defense metabolism. Directed evolution of promiscuous reactions is one of the core technologies underpinning the field of synthetic biology. Our results provide a more complete understanding of how natural selection uses promiscuous reactions to generate new enzymes in nature and chemical diversity in pathogen defense, as well as demonstrate a novel strategy for identifying their molecular origins in highly divergent, related enzymes.

## Main

Class II P450s from a monophyletic group of CYP71, CYP736 and CYP83 families have been repeatedly recruited for the biosynthesis of cyanohydrins (α-hydroxynitriles) and related structures from amino acid-derived oximes (Fig. 1a) (Bak et al., 1998; Jørgensen et al., 2011; Nafisi et al., 2007; Nelson and Werck-Reichhart, 2011; Takos et al., 2011; Irmisch et al., 2014; Rajniak et al., 2015; Yamaguchi et al., 2014, 2016). They catalyze three consecutive reactions: an E-to-Z isomerization of the oxime substrate, oxime dehydration, and hydroxylation at either the α-, β-, γ- or benzylic carbon (Bak et al., 1998; Nafisi et al., 2007; Klein et al., 2013; Clausen et al., 2015; Knoch et al., 2016). In *A. thaliana*, CYP83B1 lost the ability to isomerize tryptophan (Trp)-derived indole-3-acetaldoxime (IAOx), using instead the E-oxime as its substrate in a pathway that produces the defense metabolite 4-methoxy-indol-3yl-methylglucosinolate (4M-I3M) (Fig. 1a) (Bak et al., 2001; Bednarek et al., 2009; Clay et al., 2009; Bednarek et al., 2011; Clausen et al., 2015). By contrast, CYP71A12 and CYP71A13 - which share 89.3% sequence identity - catalyze the conversion of IAOx to form α-hydroxy-indole-3-acetonitrile (α-hydroxy-IAN), a novel cyanohydrin that is the precursor to the cyanogenic defense metabolite 4-hydroxy-indole-3-carbonylnitrile (4OH-ICN) in Arabidopsis (Fig. 1a) (Nafisi et al., 2007; Klein *et al*., 2013; Rajniak et al., 2015). CYP71A13 alone additionally dehydrates α-hydroxy-IAN to form dehydro-IAN, precursor to the non-cyanogenic defense metabolite camalexin in Arabidopsis (Klein et al., 2013).

**Fig. 1.**
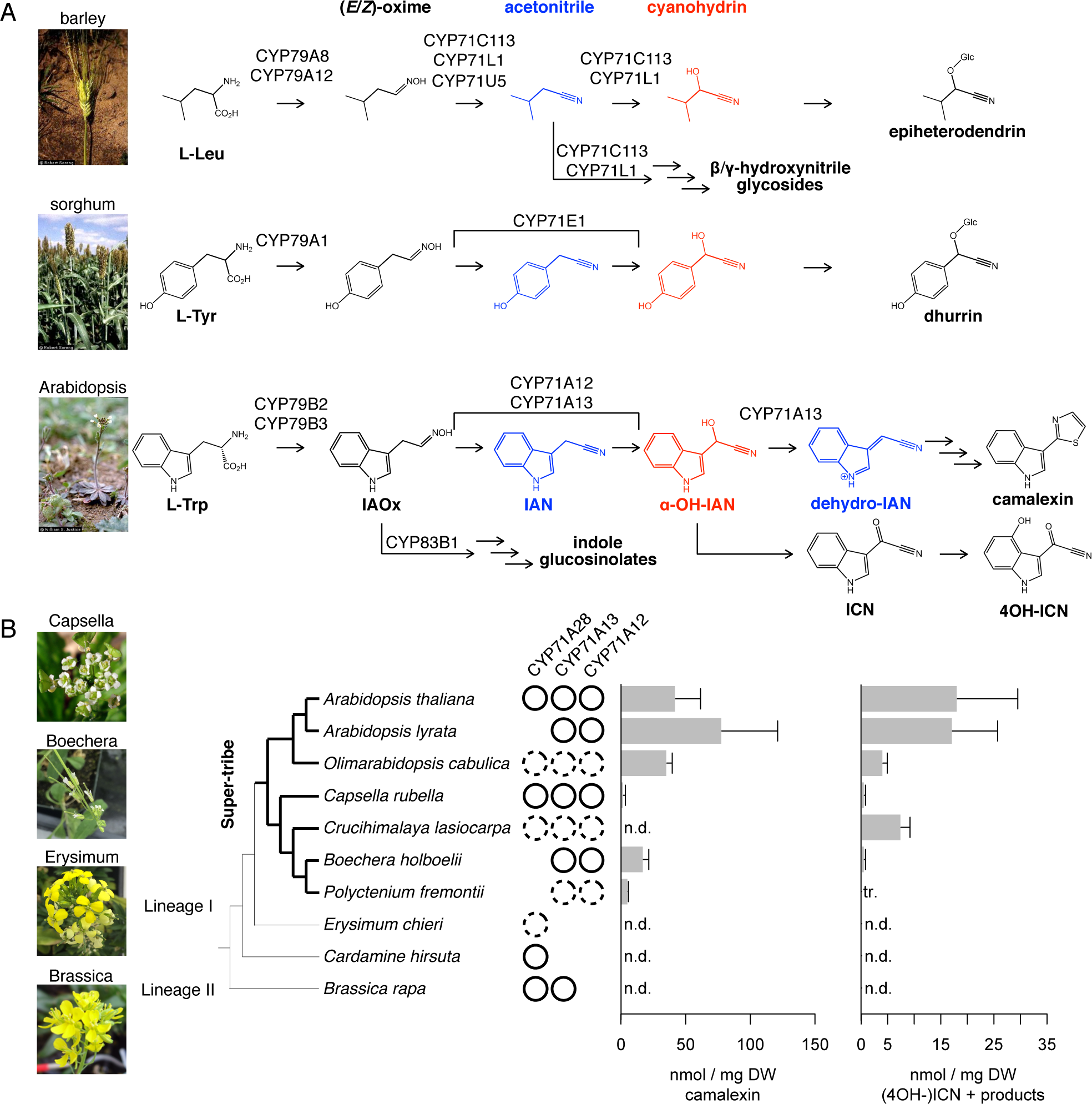
Phylogenetic distribution of CYP71A12 and CYP71A13 orthologs and their physiological functions. (a) Biosynthetic pathways of cyanohydrin-derived plant defense metabolites. Images courtesy of https://plants.usda.gov. (b) Presence of CYP71A12, CYP71A13, and CYP71A28 orthologs in the Brassicaceae, inferred from shared synteny (solid circles) or relatedness to plants with sequenced orthologs (dashed circles) See Methods for lists of contigs and chromosomes used. Phylogenetic species tree (left). Levels of camalexin (middle) and (4OH-)ICN metabolites (right) in 10-day-old seedlings inoculated with *Psta* for 30 h. Data represent mean ± SE of four replicates of 13 to 17 seedlings each. Thick branches represent the Super-tribe. To ensure robust sampling of certain lineages, unsequenced plant species were profiled, and presence of orthologs in these species were predicted based on the available sequences of close relatives. To ensure robust activation of defense metabolism in sampled species, 4M-I3M and related metabolites were profiled, and 4M-I3M’s signaling function was assayed (Extended Data Fig. 1a–b; Supplementary Note 1).

To determine whether the CYP71A13 dehydration reaction is indicative of a functional gain or loss, we investigated whether physiological functions of CYP71A12/A13 are conserved outside of Arabidopsis. To that end, we performed comparative phylogenetic and syntenic analyses to identify putative homologs/homeologs from available sequences of plants in the mustard family (Brassicaceae) (Supplementary Table 1). We then profiled (6-methoxy)camalexin and ICN metabolites in seedlings that were elicited with the bacterial pathogen *Pseudomonas syringae* pv. *tomato* DC3000 *avrRpm1* (*Psta*), using liquid chromatography coupled with diode array detection and mass spectrometry (LC-DAD-MS). The Brassicaceae is split into a larger core clade of lineage I and II species and a smaller sister clade, containing the genus Aethionema (Couvreur et al., 2009; Huang et al., 2015). CYP71A12 and CYP71A13 orthologs in a monophyletic group of lineage I species (hereafter referred to as the ‘Super-tribe’) likely contribute to the biosynthesis of (4OH-)ICN and (6-methoxy)camalexin in these species (Fig. 1b; Extended Data Fig. 1, Supplementary Note 1), whereas CYP71A13 orthologs in lineage II species do not synthesize (4OH-)ICN and (6-methoxy)camalexin (Fig. 1b; Extended Data Fig. 2d), indicating important functional differences between lineage I and II CYP71A13 enzymes.

**Fig. 2.**
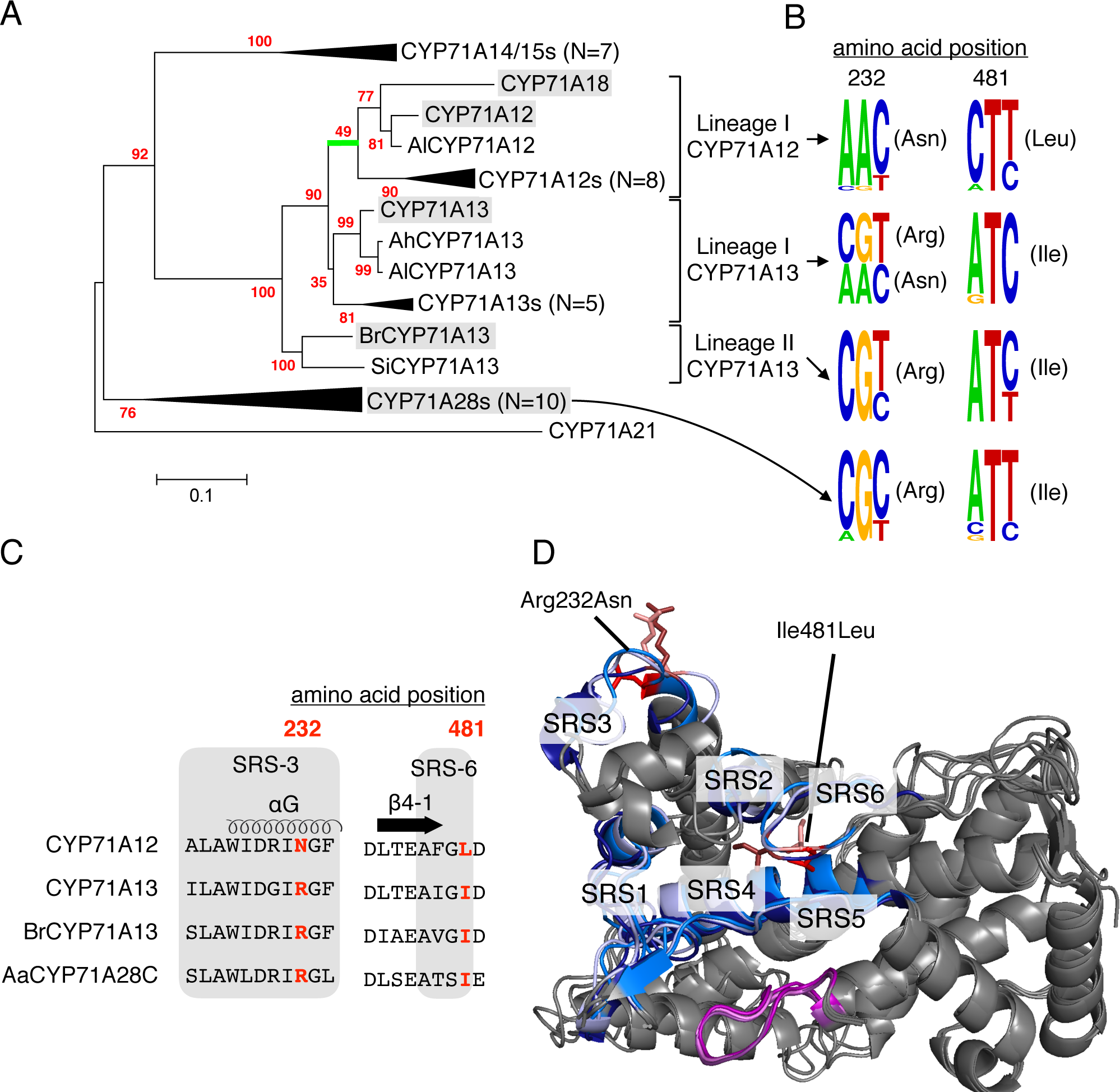
Branch-site likelihood modeltest for positive selection identifies two putative substrate-binding residues within the CYP71A12 active site. (a) Phylogenetic maximum likelihood tree of full-length CYP71A12, CYP71A13 and CYP71A28 protein sequences. Bootstrap values (N=100 replicate trees) are shown in red at the nodes. Scale bar represents 0.05 nucleotide substitutions per site. Individual branches identified to be under positive selection, using the Branch-site model (*P* < 0.05, likelihood ratio tests) are shaded in green. Proteins used in this study are boxed in grey. (b) Alignment of sequence logos of amino acid positions 232 and 481 of full-length and and partial sequences. (c) Alignment of the SRS-3 and SRS-6 sequences. Residues expected to fall within SRS sequences are boxed in gray. Residues under positive selection are in red. The conserved secondary structures are indicated at the top. (d) Structural overlay of homology models of CYP71A12 (medium-colored), and CYP71A13 (light-colored), and AaCYP71A28C (dark-colored) shown as ribbon drawings (alignment in Extended Data Fig. 3b). Colored regions indicate the locations of the six SRSs (blue) around the heme-containing active site (purple) and of the residues in positions 232 and 481 (red, in ball and stick). See Methods for lists of sequences.

CYP71A12 and CYP71A13 are paralogous enzymes that diverged from CYP71A28 through transposition and tandem gene duplications (Werck-Reichhart *et al*., 2002; Bak *et al*., 2011). Our maximum likelihood tree shows excellent support (>90%) for three distinct subclades: lineage I CYP71A12, lineage I CYP71A13, and lineage II CYP71A13 orthologs (Fig. 2a; Extended Data Fig. 2a). The absence of lineage II CYP71A12 sequences (Fig. 2a; Extended Data Fig. 2a–b) suggests either loss of CYP71A12 or non-duplication of CYP71A13 (Fig. 2a; Extended Data Fig. 2a–b). CYP71A18 is the sole nonsyntenic member of the CYP71A12/CYP71A13 clade (Fig. 2a; Extended Data Fig. 2a–b) and likely duplicated from CYP71A12 after *A. thaliana* speciation; it has no characterized function to date. The elongated branch length underlying the CYP71A12/CYP71A13 clade suggests accelerated evolution (Fig. 2a), which could be due to positive selection or neutral drift after escape from the evolutionary constraints imposed on the common ancestor. To examine the first scenario, we performed likelihood ratio tests (LRTs) based on branch-site models to detect positive selection on individual codons along prespecified lineages of our tree (Fig. 2a; Extended Data Fig. 2a) (Zhang et al., 2005). As expected, a substitution rate (*dN/dS*) below 1 (0.15-0.16) was observed for most residues throughout the tree (Extended Data Fig. 1c; Supplementary Table 1), indicating purifying selection. However, a *dN/dS* far above 1 (80.03) was observed for thirteen residues in CYP71A12 sequences against all other sequences (Fig. 2a; Extended Data Figs. 2b–c, 3a; Supplementary Table 1), indicating that these residues are conserved or under neutral selection in CYP71A13 and CYP71A28 enzymes, but are under positive selection in CYP71A12 enzymes. LRTs also accepted the branch-site model over a null hypothesis that assumes neutral evolution in the foreground lineage (*P* < 0.05; Supplementary Table 1), a more stringent test for positive selection. LRTs did not accept the branch-site model for positive selection on CYP71A18 (Supplementary Table 1), indicating CYP71A18 has escaped from the selective constraints imposed on CYP71A12.

We then investigated the natural distribution pattern of Asn232 and Leu481, which are CYP71A12 residues identified to be under positive selection with posterior probabilities ≥ 095 (Supplementary Table 1). CYP71A28 and lineage II CYP71A13 enzymes predominantly contain Arg232 and Ile481 residues (Fig. 2b). Lineage I CYP71A13 enzymes retain Ile481 but nearly half exhibit Arg232Asn substitutions (Fig. 2b), a pattern that could have been present in evolutionary intermediates between the two P450s. While most CYP71A12 enzymes exhibit Arg232Asn and Ile481Leu substitutions, CrCYP71A12L1 retains Arg232, a pattern that could represent the first mutational step in regressive evolution. An Arg232Asn substitution requires a two-base change to interconvert between Arg and Asn; we were unable to find one-base changes (leading to Ser/His) at the corresponding position in the available sequences (n = 29; see Methods), further supporting rapid CYP71A12 evolution.

The six predicted substrate recognition sites (SRS1–6) in CYP71A12s, each contains a single residue under positive selection (Fig. 2c; Extended Data Fig. 3a–b) (Gotoh 1992; da Fonseca et al., 2007; Seifert and Pleiss, 2009; Zawaira et al., 2011). Of these residues, only Asn232 and Leu481 can form a hydrogen bond and hydrophobic contacts with the anionic site and aromatic ring of the bound substrate α-hydroxy-IAN. Homology models based on six P450 crystal structures in the closed conformation predict that the Arg232Asn substitution may form a new hydrogen bond with the substrate, orienting it away from the active site (Fig. 3d). Furthermore, Ile481Leu may alter the conformation of the C-terminal loop, which houses SRS-6, to cooperatively support the exclusion of the substrate from the active site and the loss of dehydration reaction in CYP71A12 (Fig. 3d).Interestingly, a two-residue sequence gap N-terminal to SRS-6, present in CYP71A12s and CYP71A13s and not in CYP71A28s (Extended Data Fig. 3a), repositions the Ile481 sidechain so that it may form new hydrophobic interactions with the substrate (Fig. 3d) and potentiate the acquisition of dehydration reaction in CYP71A13.

**Fig. 3.**
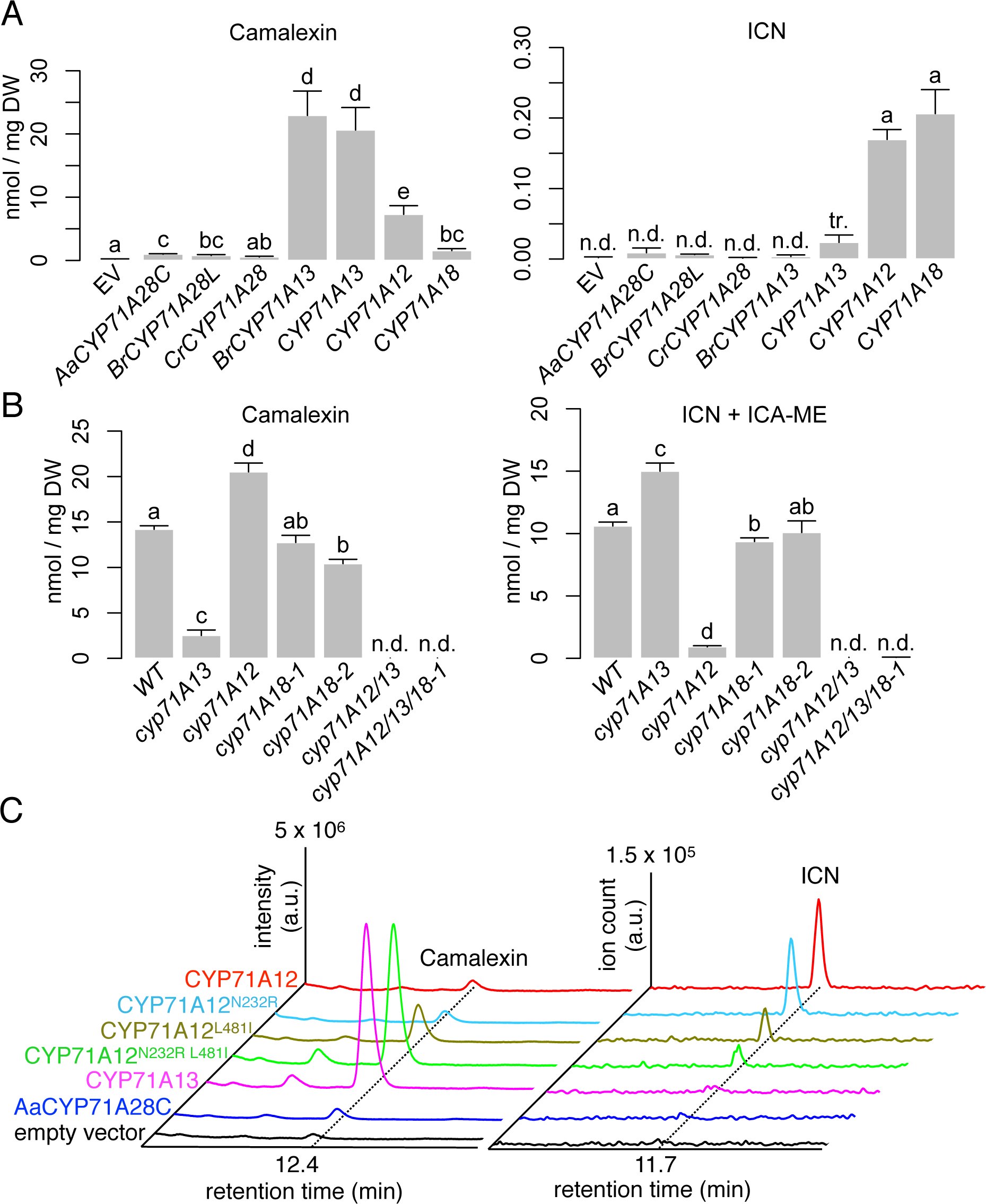
Mutagenesis and epistasis potentiated CYP71A12 functionalization. (a–b) Levels of camalexin (left) and (4OH-)ICN (right) in *N. benthamiana* leaves (a) and 10-day-old seedlings elicited with *Psta* for 24 h (b). Empty vector (EV) or indicated *CYP71* genes in (a) were co-expressed with *CYP79B2, GGP1*, and *CYP71B15* to reconstitute the camalexin pathway (left), or with *CYP79B2* and *FOX1* to reconstitute the ICN pathway (right). Data represent mean ± SE of three replicate leaves (a) and three replicates of 13-17 seedlings (b). Different letters denote statistically significant differences (*P* < 0.05, two-tailed *t* test). (c) Fluorescence emission chromatogram of camalexin (left) and extracted ion chromatogram of ICN (right) in *N. benthamiana* leaves.

To investigate whether CYP71A12 and CYP71A13 are evolutionarily constrained in their catalytic activities, we first compared the reaction product profiles of representative Brassicaceae CYP71A12, CYP71A13, and CYP71A28 enzymes by reconstituting the ICN and camalexin pathways in *Nicotiana benthamiana* and substituting enzymes at the CYP71A-catalyzed steps. ICN and camalexin production was used to infer the production of labile direct products α-hydroxy-IAN and dehydro-IAN (Klein et al., 2013). The substitution of *A. thaliana* or *B. rapa* CYP71A13 in the camalexin and ICN pathway yielded high amounts of camalexin and low or undetectable amounts of ICN, respectively, whereas the substitution of CYP71A12 or CYP71A18 yielded the near-opposite (Fig. 3a), indicating that they, like most enzymes, catalyze promiscuous reactions (Khersonsky and Tawfik, 2010; Copley, 2017). The substitution of CYP71A28 enzymes or the empty vector (EV) yielded trace and undetectable amounts of camalexin and ICN, respectively (Fig. 3a), indicating non-acquisition of specific physiological functions. The promiscuous camalexin reaction observed for EV in *N. benthamiana* is likely due to the presence of distantly-related native CYP71As (Ohkawa et al., 1998). The rise and fall in efficiencies for promiscuous secondary reactions in CYP71A13 and CYP71A12 relative to BrCYP71A13 and CYP71A18 (Fig. 3a) are likely the result of changes in selective pressures on these enzymes (Fig. 2b). Consistent with the observed promiscuous reactions *in vivo,* camalexin and (4OH-)ICN amounts are higher in *Psta*-infected loss-of-function *cyp71A12* and *cyp71A13* single mutants, respectively, than in the *cyp71A12 cyp71A13* double mutant (Fig. 3b; Extended Data Figs. 5-6, Supplementary Note 2) (Müller et al., 2015).

We next generated single and double Asn232Arg and Leu481Ile CYP71A12 revertants and compared their reaction profiles to CYP71A12 and CYP71A13. The CYP71A12 Asn232Arg reversion did not alter catalytic specificities or efficiencies (Fig. 3c; Extended Data Fig. 4b). The Leu481Ile reversion yielded intermediate amounts of both ICN and camalexin (Fig. 3c; Extended Data Fig. 4b).This previously undocumented catalytic activity may have been present in evolutionary intermediates; confirming this, we observed a Leu481Ile substitution in at least one CYP71A12 ortholog (Fig. 2b). The CYP71A12 double revertant enzyme completely inverted the catalytic specificities to that of CYP71A13 and notably restored the dehydration reaction to dehydro-IAN (Fig. 3c; Extended Data Fig. 4b). These results indicate that the catalytic specificities and efficiencies between CYP71A12 and CYP71A13 are mutationally accessible, requiring as little as three base changes. Furthermore, the regressive evolution of CYP71A12 involves an epistatic interaction on successive mutations; whereby the Asn232Arg and Leu481Ile substitutions respectively potentiate and actualize the ability to biosynthesize dehydro-IAN.

**Fig. 4.**
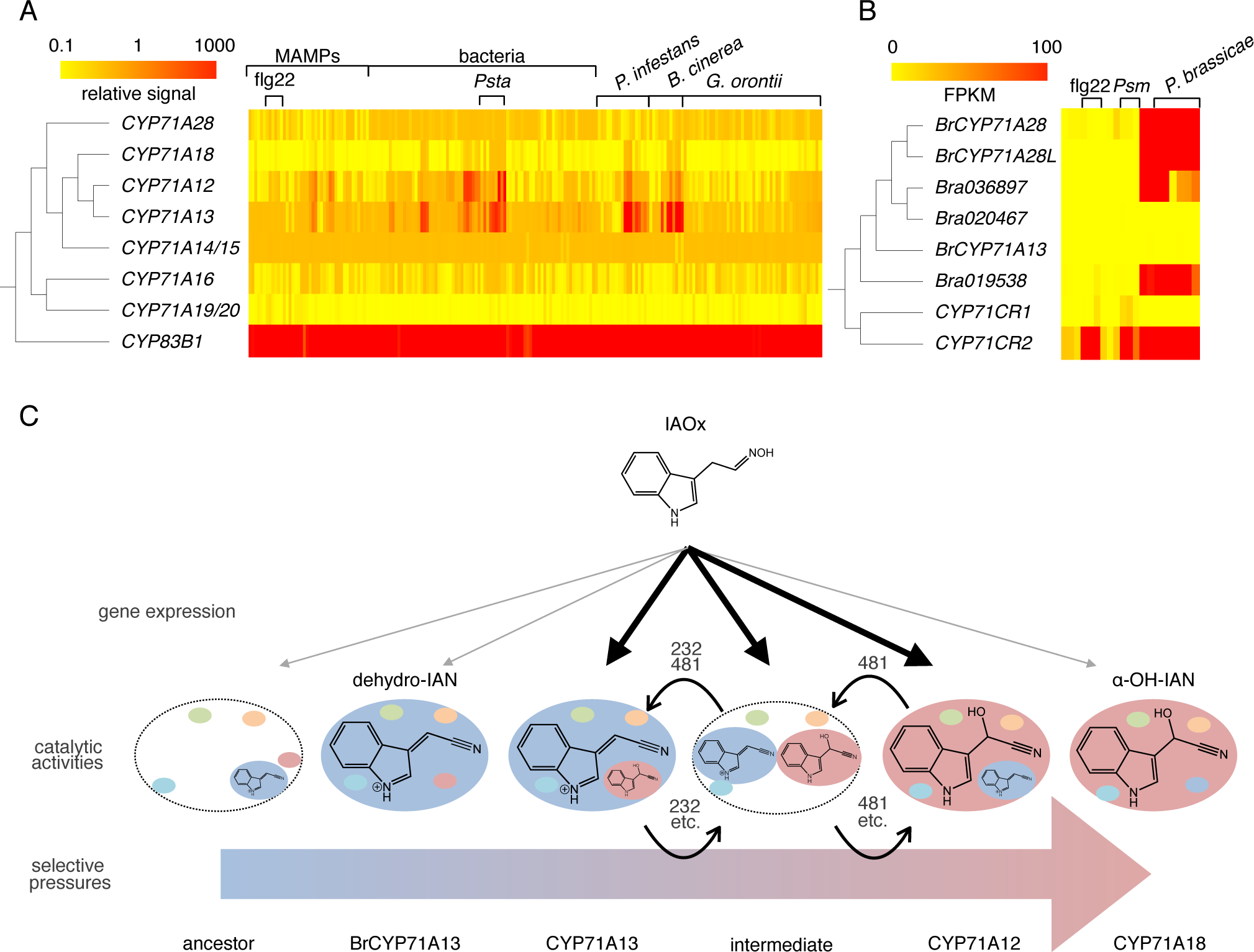
Changes in gene expression potentiated neofunctionalization of CYP71A13 and release of CYP71A18 from selective constraint. (a) Heat map of gene expression for P450 genes in Arabidopsis under various pathogen treatments. (b) Heat map of gene expression for P450 genes in B. rapa under two pathogen treatments. (c) Proposed evolutionary trajectory of CYP71A enzymes involved in cyanogenic defense metabolism in Arabidopsis.

In the course of this study, we observed that the primary reactions of BrCYP71A13 and CYP71A18, while apparent via heterologous expression (Fig. 3a), could not be detected in their *Psta*-infected native hosts (Figs. 1b and 3b). These results suggest that BrCYP71A13 and CYP71A18, which book-end the evolutionary trajectories of CYP71A13 and CYP71A12, are not natively expressed in a pathogen-inducible manner. To test this hypothesis, we obtained datasets for all transcriptomics experiments on biotic stress in Arabidopsis and *B. rapa* (Toufighi et al., 2005; Klein et al., 2015; Chen et al., 2016), and examined expression patterns of all full-length CYP71A members. CYP71A18’s gene expression is mostly unchanged between mock and pathogen treatments and at least 22 to 467-fold less than those of CYP71A12/CYP71A13/CYP83B1 for a variety of pathogen treatments (Fig. 4a). Similarly, BrCYP71A13’s gene expression is 44 to 9185-fold less than those of related CYP71CRs in *B. rapa* defense metabolism (Fig. 4b) (Klein et al., 2015; Chen et al., 2016). Altogether, our results suggest that changes in gene expression and substrate-recognition sites were both necessary to increase the efficiency and thus the physiological relevance of catalytic promiscuity in CYP71A13 and CYP71A12, respectively (Fig. 4c).

Proteins evolve by following trajectories through ‘rugged’ sequence space, consisting of mutational steps between functional proteins and between their catalytic specificities (Smith, 1970; Kondrashov & Kondrashov, 2015; Starr and Thornton, 2016). The rise in catalytic promiscuity is likely responsible for the repeated evolution of cyanohydrins and their cyanogenic and non-cyanogenic derivatives as defense chemicals in over 2,650 higher plant species and certain arthropods, fungi and bacteria (Nahrstedt 1985; Davis and Nahrstedt 1987; Blumer and Haas, 2000; Zagrobelny et al., 2007, 2008; Klein et al., 2013; Caspar and Spiteller, 2015; Rajniak et al., 2015). We observed that promoter mutations and intramolecular epistasis rendered accessible the evolutionary trajectories from CYP71A12 to CYP71A13, BrCYP71A13 to CYP71A13, and CYP71A12 to CYP71A18, and the resulting increases and decreases in catalytic promiscuity. Collectively, our data provide additional insight into the evolvability of promiscuous activities for the discovery and directed evolution of novel enzymes.

## METHODS

Information on Abbreviations, Plant Materials and Growth Conditions, Pathogen infection, Extraction and LC-DAD-MS Analysis of (6-methoxy)camalexin and (4OH-)ICN, Extraction and LC-DAD-FLD-MS Analysis of Glucosinolates, Identification of *CYP71A13* and *CYP71A12* orthologs, 4M-I3M Activity Assay and Identification of callose pathway orthologs, and Heterologous pathway reconstitution in *N. benthamiana* can be found in Supplementary information.

## SUPPLEMENTARY INFORMATION

Additional supplementary information and extended data are available for this paper.

### ACKNOWLEDGEMENTS

We thank JL Celenza for *cyp79B2 cyp79B3*, E Glawischnig for (*cyp71A12 cyp71A13)-1*, and T Mitchell-Olds for *Boechera stricta* SAD12/LTM and *Boechera holboelii* 910/913. We thank ES Sattely for (4OH-)ICN and camalexin standards. This work was supported by T32-GM007499 (to BB) and Elsevier/Phytochemistry Young Investigator Award (to NKC).

## AUTHOR CONTRIBUTIONS

BB and NKC performed 4M-I3M activity assays. BB and LZ performed indole glucosinolate profiling. BB performed all other experiments. BB and NKC interpreted the results and wrote the paper.

## AUTHOR INFORMATION

The authors declare no competing financial interests. Correspondence should be addressed to B. Barco

